# White matter connectivity between occipital and temporal regions involved in face and voice processing in hearing and early deaf individuals

**DOI:** 10.1101/339713

**Authors:** Stefania Benetti, Lisa Novello, Chiara Maffei, Giuseppe Rabini, Jorge Jovicich, Olivier Collignon

**Affiliations:** Center for Mind/Brain Studies, University of Trento, 38123 Trento, Italy; Athinoula A. Martinos Center, Massachusetts General Hospital, Charlestown, MA 01129, USA; Institute of Research in Psychology (IPSY) and in Neuroscience (IoNS), University of Louvain, 1348 Louvain-la-Neuve, Belgium

**Keywords:** Cross-modal plasticity, Anatomical-functional connectivity, Deafness, Diffusion-based tractography, Temporal voice area

## Abstract

Neuroplasticity following sensory deprivation has long inspired neuroscience research in the quest of understanding how sensory experience and genetics interact in developing the brain functional and structural architecture. Many studies have shown that sensory deprivation can lead to cross-modal functional recruitment of sensory deprived cortices. Little is known however about how structural reorganization may support these functional changes. In this study, we examined early deaf, hearing signer and hearing non-signer individuals using diffusion MRI to evaluate the potential structural connectivity linked to the functional recruitment of the temporal voice area by face stimuli in deaf individuals. More specifically, we characterized the structural connectivity between occipital, fusiform and temporal regions typically supporting voice- and face-selective processing. Despite the extensive functional reorganization for face processing in the temporal cortex of the deaf, macroscopic properties of these connections did not differ across groups. However, both occipito- and fusiform-temporal connections showed significant microstructural changes between groups (fractional anisotropy reduction, radial diffusivity increase). We propose that the reorganization of temporal regions after early auditory deprivation builds on intrinsic and mainly preserved anatomical connectivity between functionally specific temporal and occipital regions.

**Highlights:**

- Macrostructural connectivity of the face-voice system is preserved in early deafness
- Early deafness impacts on the microstructural connectivity of the face-voice system
- Both genetics and experience shape structural connections in the face-voice system
- Innate anatomical networks might constrain the expression of cross-modal plasticity

**Abbreviations:**

STG: Superior Temporal Gyrus
STS: Superior Temporal Sulcus
pSTS: Posterior superior Temporal Sulcus
FFA: Face-fusiform Area
V2/3: Boundary between Extrastriate V2 and V3 Areas
TVA: Temporal Voice-sensitive Area

## Introduction

Decades of neuroscientific research have revealed the extraordinary capacity of the human brain to adapt in response to experience and lack of specific sensory inputs (Pascual-Leone et al. 2005). After sensory deprivation, such as blindness or deafness, the sensory deprived cortices can reorganize and process information from the spared sensory modalities (Bavelier and Neville 2002; Heimler et al. 2014). In the case of deafness, for instance, temporal auditory regions can be functionally recruited to respond to visual (Finney et al. 2001; Fine et al. 2005) and tactile (Auer et al. 2007; Karns et al. 2012) inputs.

It has been suggested that cross-modal reorganization following early sensory deprivation reflects the functional specialization of the colonized cortical regions (Dormal and Collignon 2011; Reich et al. 2011; Ricciardi et al. 2014). For instance, Lomber and colleagues have reported that, in deaf cats, superior visual motion detection is selectively impaired if a specific region in the dorsal auditory cortex, which processes auditory motion in hearing cats, is transiently suppressed (Lomber et al. 2010). In deaf humans, supporting evidence has recently been provided by a study showing rhythm-specific visual activations in posterior-lateral and associative auditory regions (Bola et al. 2017) and, further, by our observation of preferential responses to faces and face discrimination in the human temporal voice sensitive area (TVA) as a consequence of early auditory deprivation (Benetti et al. 2017).

How does specific non-auditory information reach the reorganized temporal cortex of deaf individuals? Evidence that cross-modal remapping of temporal regions is associated with reorganization of long-range functional interactions between auditory and visual cortices has been reported for visual motion detection in deafness (Shiell et al. 2014). Further, in our previous study we reported that face-selective cross-modal activation in the deaf TVA is primarily modulated by increased feed-forward effective connectivity from extrastriate visual regions (V2/3) in early deaf humans (Benetti et al. 2017). This observation confirms previous findings reported in early blind individuals (Collignon et al. 2013) and suggests that the reorganization of long-range functional connectivity between sensory cortices might play a key role in functionally selective cross-modal plasticity.

Despite the growing evidence of selective neurofunctional plasticity in both blindness and deafness, whether these changes relate to alterations of white matter structural connections still remains controversial. In particular, the observation of increased functional connectivity within reorganized networks (e.g. between temporal and occipital regions) seems not to be systematically paralleled by consistent observations in structural connectivity, where both reductions and increases have been observed in blind (Shu et al. 2009; Lao et al. 2015; Bauer et al. 2017) and deaf (Lyness et al. 2014; Karns et al. 2017) individuals. This inconsistency might be due to the fact that previous studies have mostly focused on functional and structural connectivity separately.

Reorganization of the face-voice human system in early deafness represents a unique opportunity to specifically address the relationship between changes in local cortical responses and long-range functional and structural connectivity within a functionally defined network. In fact, there is evidence of direct connections between the fusiform face-selective area (FFA) and the mid-anterior portion of the superior temporal gyrus (STG) responding selectively to human voices in the right hemisphere of hearing individuals, (i.e. TVA; Blank et al. 2011) as well as direct connections between extrastriate visual and temporal auditory regions in humans (Beer et al. 2011).

In this study, we follow-up on our previous observation (Benetti et al. 2017) by applying a hypothesis-driven approach to the examination of white matter connectivity in the same deaf individuals showing face-selective reorganization of both temporal regions and long-range occipitotemporal functional coupling.

## Materials and Methods

### Participants

Forty-four participants were included in this study. Fourteen were early deaf (ED; 13 since birth and one before age 4; mean age 32.79 ± 7.21; 7 males), 15 were hearing controls fluent in the use of the Italian Sign Language (HC-SL; mean age 34.07 ± 5.97; 5 males) and 15 were hearing controls (HC; mean age 30.40 ± 5.09; 8 males). All the participants participated also in our study on face-selective cross-modal plasticity in the temporal voice area (Benetti et al. 2017). ED participants presented with severe to profound congenital deafness in both ears with the exception of one participant who developed deafness before the age of 4 years. A group of hearing sign language (Lingua Italiana dei Segni; LIS) users was included to control for the potential confound of visual language use; the age of acquisition and the exposure to LIS use was comparable between the ED and HC-SL groups (see Table 1). All participants were currently healthy, had no medical history of neurological or psychiatric disorders and were no under psychoactive medication. They were tested on non-verbal IQ, hand preference and composite measures of face recognition (a detailed description of these tests has been provided in Benetti et al. 2017). There were no differences for age, non-verbal IQ scores and hand preferences between groups while both the ED and HC-SL outperformed the HC group on face recognition (Table 1). The study was approved by the Committee for Research Ethics of the University of Trento; all participants gave informed consent in agreement with the ethical principles for medical research involving human subject (Declaration of Helsinki, World Medical Association) and the Italian Law on individual privacy (D.1. 196/2003).

**Table 1.** Demographies, behavioral performances and Italian Sign Language aspects of the 44 subjects participating in the diffusion and functional MRI experiment

| Demographics/Cognition | Participants in Diffusion/Functional MRI Experiment |  |  | Statistics |
| --- | --- | --- | --- | --- |
|  | Hearing Non-Signers (n =15) | Hearing Signers (n=14) | Early Deaf (n=14) |  |
| Mean Age, y(SD) | 30.40 (5.09) | 34.07(5.97) | 32.79(7.21) | F-test=1.382<br>P-value = 0.263 |
| Gender, male/female | 8/7 | 5/10 | 7/7 | X<sup>2</sup>= 1.381<br>P-value = 0.501 |
| Hand Preference, % (Right/Left ratio) | 71.36(48.12) | 61.88(50.82) | 58.63(52.09) | K-W test = 1.15<br>P-value = 0.564 |
| Non-verbal IQ, Mean estimate (SD) | 123.4(8.38) | 124.80(5.61) | 120.23(9.71) | F-test=1.073<br>P-value = 0.352 |
| Composite Face Recognition, z-scores (SD) | -0.124(1.45)** | 1.547(1.43) | 1.634(1.74) | F-test=5.751<br>P-value = 0.006 |
| LIS Exposure, y (SD) | -- | 25.03(13.85) | 21.08(10.25) | t-test=-0.823<br>P-value =0.418 |
| LIS Acquisition, y (SD) | -- | 11.38(9) | 9.033(11.91) | M-W.U-test = 112<br>P-value =0.505 |
| LIS Frequency, Percent time/y (SD) | -- | 68.59(44.88) | 84.80(26.09) | M-W. U-test = 87.5<br>P-value=0.364 |

(\*\*) Main effect of group ( $P < 0.006$ ), ED versus HC ( $p < 0.016$ ), HS-SL versus HC ( $p < 0.016$ ). K-W, Kruskal-Wallis; M-W, Mann-Whitney; y, years; SD, Standard Deviation.

### Image acquisition

Four imaging datasets were acquired for completion of this study: a functional MRI face localizer, a functional MRI voice localizer, diffusion-weighted MR images and structural T1-weighted images. Data acquisition was carried out at the Center for Mind/Brain Science of the University of Trento, using a Bruker BioSpin MedSpec 4T MR-scanner.

#### Functional MRI sequence and experiment

A detailed description of the experimental procedure and stimuli for the face and voice localizers have been provided elsewhere (Benetti et al. 2017). In brief, the same gradient-echo planar imaging (EPI) sequences with continuous slice acquisition (TR=2.200 ms, TE = 33ms, slices = 37, slice thickness = 3mm, Slice gap = 0.6mm, image matrix = 64 × 64) was applied and a total of 274 and 335 volumes were acquired for the face and voice localizer respectively. The voice localizer was acquired in the HC group only during a 12 minutes task in which neutral human vocal sound (“A”), scrambled human vocal (random mix of magnitude and phase of each Fourier component but same global energy and envelope of neutral vocal stimuli) and object sounds were presented in a block-design manner (10 blocks for each category). The face localizer also consisted of a block-design task in which images of human faces (static and neutral) and two-floor houses were alternated across 20 blocks (10 for each category, 20 images for block) for a total duration of about 10 minutes. All the images had been previously equated for low-level properties such as mean luminance, contrast and spatial frequencies (Shine Toolbox; Willenbockel et al. 2010). The selection of the 4 seed regions used for white matter tracking was implemented by contrasting the face/house and human vocal/object sounds within each experiment as described in the dedicated section below (*Definition of regions of interest (ROIs) for tractography)*.

#### Diffusion MRI (dMRI)

Whole brain diffusion weighted images were acquired using an EPI sequence (TR = 7100 ms, TE = 99 ms, image matrix = 112 × 112, FOV = 100 × 100 mm^2^) with a b-value of 1500 s/mm^2^. Ten volumes without any diffusion weighting (b0-images) and 60 diffusion-weighted volumes (slice thickness = 2.29 mm) were acquired. By using a large number of directions and ten repetitions of the baseline images on a high magnetic field strength we aimed at improving the signal-to-noise ratio and reduce implausible tracking results (Fillard et al. 2011).

#### T1 images

To provide detailed anatomy a total of 176 axial slices were acquired with a T1-weighted MP-Rage sequence covering the whole brain. The imaging parameters were: TR = 2.700 ms, TE= 4.18ms, flip angle = 7°, isotropic voxel = 1 mm^3^.

### Image processing

#### dMRI preprocessing

Raw images were visually inspected for artifacts and a quality check was also carried out using the Data Quality Tool (Version 1.5, 2014, New York University Centre for Brain Imaging, http://cbi.nyu.edu/software/dataQuality.php). Spatial and temporal statistics were extracted from a number of ROIs inside and outside the brain before computing individual mean Signal-to-Noise ratio (SNR) and number of spikes (for details on how SNR was computed see Velasco P., 2006, Technical Report at http://cbi.nyu.edu/Downloads/dataQuality.pdf). Participants were included if they presented with less than 3 spikes across all volumes; additionally, according to our DWI sequence parameters, we set the inclusion criteria for mean individual SNR to >10 (Choi et al. 2011). Based on these criteria, no participants were excluded from the study: the number of spikes did not differ from zero within groups (HC = 0.0 ± 0.0, HC-SL = 0.06 ± 0.258, ED = 0.0 ± 0.0) and no significant differences were detected for mean SNR across groups (HC = 13.27 ± 0.356, HC-SL = 13.05 ± 0.356, ED = 12.95 ± 0.369; *F-test* = 0.210, *p-value* = 0.812).

Data pre-processing was implemented in FSL 5.0.9 (http://fsl.fmrib.ox.ac.uk/fssl/fslwiki). Data were corrected for Eddy currents distortions and head motion by registering the DW-volumes to the first b0 volume. The gradient direction (b-vec) for each volume was corrected applying the individual rotation parameters (Leemans and Jones 2009) and non-brain voxels were removed with FSL-BET. Subsequently, the diffusion tensor model was fitted to the data and the fractional anisotropy (FA), radial diffusivity (RD) and axial diffusivity (AD) indices computed for each voxel in the brain (Basser and Pierpaoli 2011).

In each voxel, from the measured data, local fiber bundle orientations were estimated by using FSL-BEDPOSTX and applying a ball-and-stick model (Behrens et al. 2007) which allowed to estimate a maximum of two-fiber directions in each voxel. Finally, a series of registration matrices were calculated for transforming between native diffusion, native structural and standardized MNI space. This was necessary in order to transform the seed and target regions from functional MNI space to diffusion space for probabilistic fiber tractography (see preparation of seed and target regions below).

#### Definition of regions of interest (ROIs) for tractography

The pre-processing and analysis of functional data have been detailed elsewhere (Benetti et al. 2017). For the purpose of this study, four seed regions were selected based on previous studies of face-voice integration in the brain (Blank et al. 2011, 2015) and our observations on regional responses and functional/effective connectivity: FFA (Kanwisher et al. 1997; Rossion et al. 2012) in the right mid-fusiform gyrus and V2/3 in the right infero-middle occipital gyri, which showed enhanced functional coupling with the reorganized TVA; right TVA (Belin et al. 2000) in the right mid-superior temporal sulcus as voice-selective region showing cross-modal reorganization; and right posterior superior temporal sulcus (pSTS) as exclusion mask region, since it shows multimodal face responsiveness in hearing individuals and we aimed at exploring ‘direct’ V2/3- and FFA-TVA connections by excluding those fibers passing through the multimodal associative relay in pSTS. For each group, and based on the functional face-localizer experiment, the localization of face-sensitive FFA and pSTS was defined on the peak-coordinate of the group-maxima for the contrast [face > houses] while the localization of V2/3 was defined on the peak-coordinate of the contrast-maxima showing increased face-specific functional coupling with the reorganized TVA in the right hemisphere of ED compared to both HC and HC-SL groups (Benetti et al. 2017). For both the HC and HC-SL participants, the localization of voice-sensitive area TVA was defined on the peak-coordinate of the HC group-maxima for the contrast [(human voice > object sound) n (human voice > scrambled voice)] as revealed by the functional voice-localizer experiment. In ED individuals, the reorganized TVA was defined on the peak-coordinate of the group-maxima for face-selective cross-modal response in the mid-superior temporal sulcus as revealed by the face-localizer experiment. Since we applied a strong hypothesis-driven approach in this study, we restricted the focus of investigation on the right hemisphere where face-selective functional reorganization was specifically observed in the ED group. The MNI peak-coordinates for each group and each region are reported in supplemental Table 1.

**Supplemental Table 1.** Group-specific pcak-coordinatcs (MNI) used to define seed and target regions in probabilistic fiber tracking.

| Area | X<sub>(mm)</sub> | Y<sub>(mm)</sub> | Z<sub>(mm)</sub> |
| --- | --- | --- | --- |
| Right TVA in HC and HC-SL group | 63 | -22 | 4 |
| Right reorganized TVA in ED | 62 | -18 | 2 |
| Right FFA in ED | 48 | -56 | -18 |
| Right FFA in HC | 44 | -50 | -16 |
| Right FFA in HS | 44 | -52 | -18 |
| Right pSTS in ED | 50 | -44 | 14 |
| Right pSTS in HC | 52 | -42 | -16 |
| Right pSTS in HS | 52 | -44 | 10 |
| Right V2/3 in ED | 26 | -94 | 4 |
| Right V2/3 in HC | 28 | -86 | 4 |
| Right V2/3 in ED | 27 | -92 | -1 |

HC, Hearing Control; HC-SL, Hearing Signers; ED, Early Deaf; TVA, Temporal Voice-selective Area; FFA, Fusiform Face Area; pSTS, posterior Superior Temporal Sulcus; V2/3, Extrastriate V2-V3 boundary.

#### Preparation of ROIs for probabilistic tractography

First, a sphere of 8mm radius was centered at the group peak-coordinates (selected as detailed in the previous section) for each seed and target region. The reconstruction of white matter connectivity between functionally defined regions, that are likely to encompass portions of grey matter, can be challenging (Whittingstall et al. 2014). Accordingly, each group-specific sphere was moved into standard MNI-structural space (MNI-152 template) and a custom algorithm was applied that further moved them to the nearest two contiguous white matter voxels. In addition, to ensure tracking only from white matter voxels and to maintain anatomical proximity we masked seed and target regions both with a white matter skeleton mask (Blank et al. 2011) and an anatomical mask of right STG for TVA/pSTS, of right fusiform gyrus for FFA and of right inferior and middle occipital gyri for V2/3. The MNI-diffusion registration matrices were finally applied to each region to moved them into native diffusion space of each individual, depending on the belonging group. This resulted in a different number of voxels between seed/target regions and between groups, we therefore accounted for these differences in the subsequent analyses.

#### Probabilistic Tractography

For each participant, probabilistic tractography was computed by applying FSL-PROBTRACKX (Behrens et al. 2007) in symmetric mode (i.e. seeding respectively from one ROI and targeting the other in each connection) to detect right V2/3-TVA and right FFA-TVA connections. This resulted in an estimate of the most likely pathway connecting each pair of ROIs, namely the probability of connection measured as the number of streamlines successfully reaching a target voxel from a given seed. A standard parameter setting that was used for fiber tracking in this study: 5000 sample tracts per seed voxel, a curvature threshold of 0.2 (bending angle no smaller than 80°), a step length of 0.5 mm and a maximum number of 2000 steps. This parameter profile was successfully applied in a previous study investigating the white matter connection between voice and face selective regions in hearing individuals (Blank et al. 2011). To aim at connection specificity and for better anatomical precision, the white matter mask for right pSTS was used as exclusion mask (see also ROIs definition section) to exclude fibers passing through multimodal relays in the reconstruction of right V2/3-TVA and FFA-TVA connections.

### Data analysis

#### Macrostructural connectivity measures

The probability of connection for each pair of regions was determined on the number of streamlines reconstructed (sum of both directions) between them. It must be noted that currently no agreement on statistical thresholding of probabilistic tractography exists. Moreover, the detection of connections with high curvature along the pathway can be challenging in tractography and can be influenced by individual variations in cortical gyrifications and anatomy. Accordingly, the operational criterion for defining a connection as reliable at the individual level was set to a minimum number of 10 streamlines between each pair of regions. This criterion has been previously applied to reduce false-positive connections while preserving true connection sensitivity in studies segregating white matter bundles related to language (Makuuchi et al. 2009) and face-voice recognition (Blank et al. 2011).

Based on previous observations (Saur et al. 2008; Doron et al. 2010; Blank et al. 2011), we defined a threshold for connection reliability at the group level as a minimum of 50% of participants presenting the connection in each experimental group. In addition, we computed a quantitative measure of strength connectivity index (Eickhoff et al. 2010; Blank et al. 2011) that was defined as the ratio between the number of streamlines between each pair of regions and the total number of streamlines for all seed/target regions, normalized by the number of voxels in each pair to account for differences in size of seed and target regions.

#### Microstructural connectivity measures

In addition to macrostructural measures we also extracted tensor-derived quantitative measures of microstructural diffusivity for each connection (i.e. FA, RD, AD see data pre-processing section above). Reconstructed pathways were binarized and used as region of interest to extract mean FA, RD and AD values at the individual level. To further exclude possible false-positive components in each connection, the individual number of streamlines per tract voxels were divided by the total number of individual tract streamlines before binarizing them at a threshold of 0.05 (Javad et al. 2014), hence excluding all voxels with a low probability of belonging to the connection.

#### Statistical analysis

The main dependent measures in this study were computed for the two connections V2/3-TVA and FFA-TVA in the right hemisphere and were: the number of participants presenting each connection according to the operational criterion described above, the connectivity and the microstructural indices for each connection. For macrostructural measures, the presence of a connection across groups was assessed with two binomial tests while group differences in the number of subjects presenting each tract was assessed with a series of Pearson’s chi-square tests; for both binomial and Pearson’s chi-square tests significance was thresholded at *p* = 0.05 Bonferroni-corrected for multiple comparisons. Since connectivity indices were not normally distributed (Shapiro-Wilk test), we used non-parametric tests to examine differences between connections (Wilcoxon-Paris Test) and between groups (Kruskal-Wallis Test).

The assessment of the microstructural indices was carried out by implementing mixed-design ANCOVAs that included Group (HC, HC-SL, ED) as a between subject factor, Connection (V2/3-TVA, FFA-TVA) as a within subject factor and size (in voxels) of seed/target regions as a variable of no interest to exclude potential confounding effects. Holm-Bonferroni (HB) correction was applied to adjust the significance values (Holm 1979; Abdi 2010). Finally, for the microstructural measures that differed between groups, we carried out a series of bivariate correlations to explored their relationship with individual face-selective responses in right TVA in the same ED group (Benetti et al. 2017), as well as with behavioral measures of face recognition across the three groups and in the ED group alone. The assumption of distribution normality was assessed prior to correlation and Pearson’s or Spearman’s coefficients applied accordingly. The statistical threshold for significance was again corrected by applying the Holm-Bonferroni correction since the measures we correlated were unlikely to be independent of each other.

## Results

### Face selectivity in the ‘deaf’ temporal voice selective area

In our previous work (Benetti et al., 2017) we reported that deaf individuals preferentially activate a specific and discrete portion of the TVA, overlapping with a region showing functional preference for voices in hearing people. Further, this reorganized region is capable of face identity processing and selectively responds to faces at similar timing as typical face sensitive regions in the visual cortex. We also showed that increased feed-forward effective connectivity from early visual V2/3 regions primarily sustain the face selective response detected in the ‘deaf’ TVA, suggesting a functional reorganization of long-range connectiivity within the face-voice brain network associated with congenital or early deafness.

### White matter macrostructure between V2/3-TVA and FFA-TVA connections

Based on the operational criterion described in the methods section, we found evidence for structural connections between right TVA and both FFA and V2/3 in the hearing and deaf groups (Fig. 1 and Fig. 2). This was confirmed when the three groups were merged together and presence of a connection across groups was statistically assessed with binomial tests: V2/3-TVA presence proportion = 0.864, *p* = 0.001; FFA-TVA presence proportion = 0.727, *p* = 0.002. The V2/3-TVA and FFA-TVA structural connections were characterized and compared in terms of both ratio of subjects having them and connectivity strength indices (Table 2a and Fig. 3). In terms of subject ratio, Pearson’s chi-square tests showed that the number of subjects presenting the V2/3-TVA and FFA-TVA connections did not differ significantly between groups (*X^2^*= 1.14, *p* = 0.566; *X^2^*= 3.18, *p* = 0.203). Finally, within the ED group only, a weak trend for higher subject ratio was detected for the V2/3-TVA connection compared to the FFA-TVA one (*X^2^* = 2.80, sig. = 0.094).

**Figure 1.**
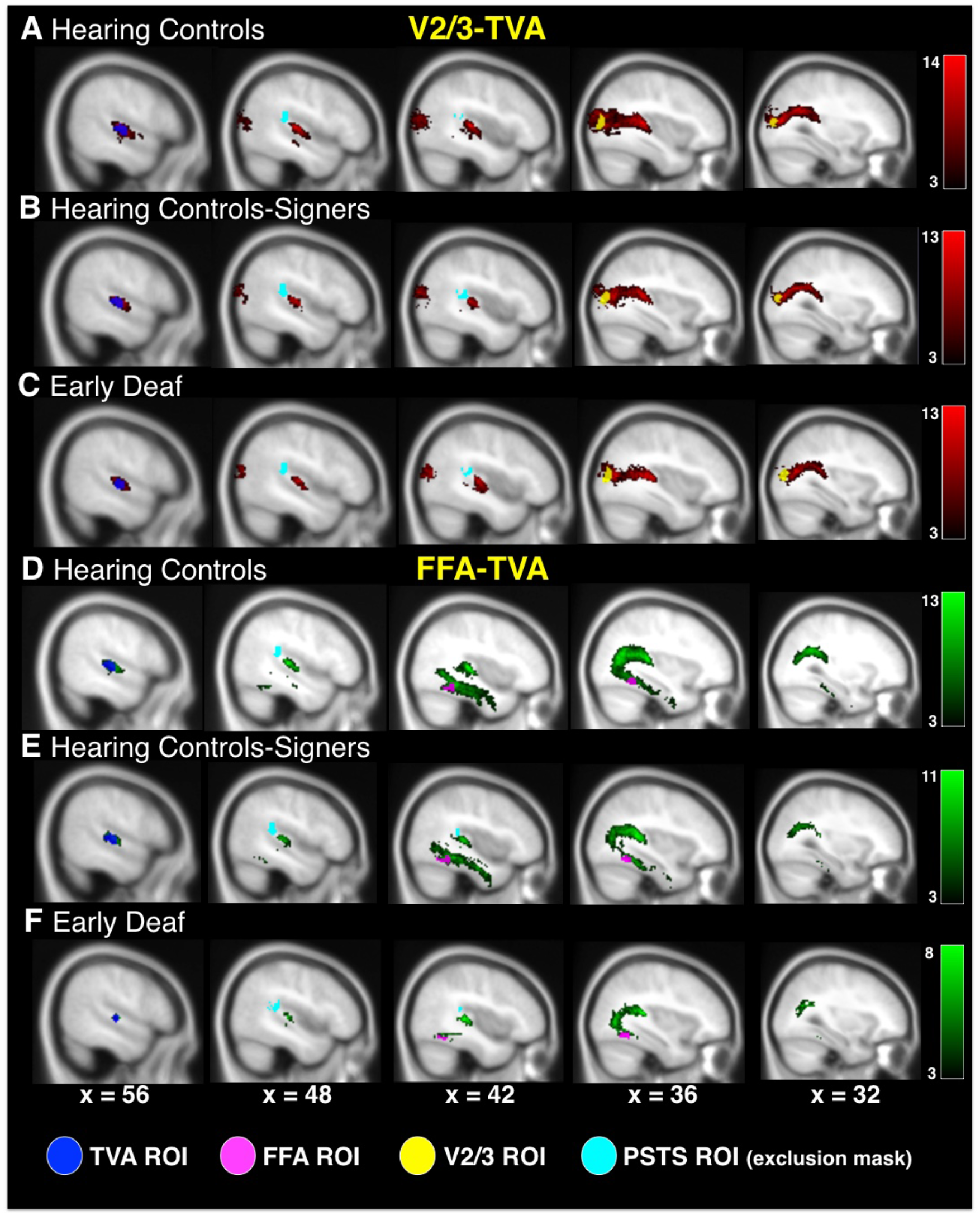
Group-specific overlays of probabilistic pathways between face-responsive V2/3 and FFA, and voice sensitive TVA. For each group, individual dMRI connectivity distributions, first thresholded at 10 paths per voxel, were then binarized and overlaid for display purposes. (**A-C**), tracking results for the V2/3-TVA path are reported in red; (**D-F**), tracking results for the FFA-TVA path are reported in green. Seed, target and exclusion regions are reported in plain colors. Bar-color codes the number of subjects showing the path in each voxel (thresholded at > 3 subjects). Tracking results are depicted on the T1 MNI-152 template.

**Figure 2.**
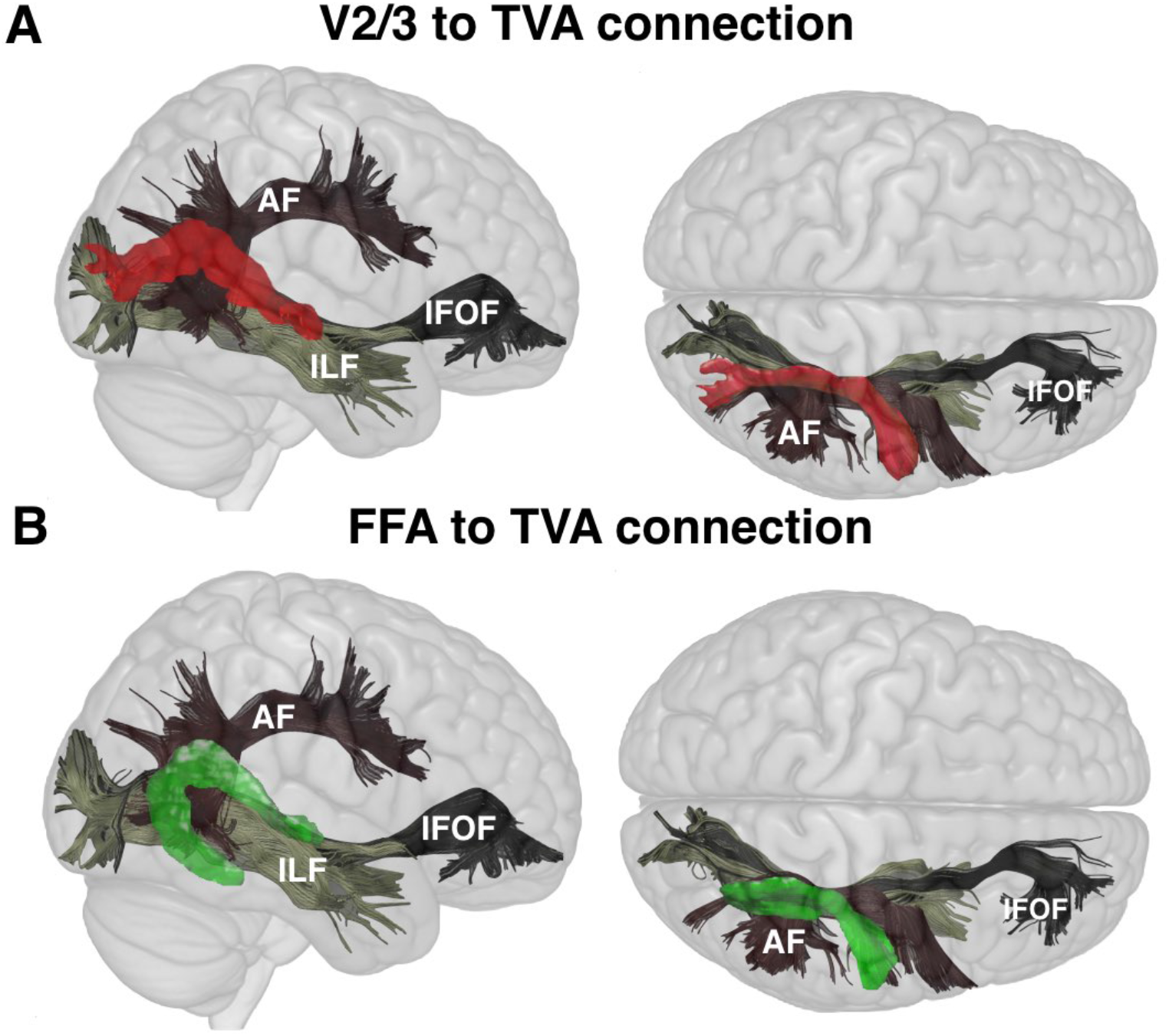
Structural pathways between face-sensitive (FFA) and voice-selective (TVA) areas as reconstructed with probabilistic fiber tracking. Results from the Hearing Control (HC) group are shown as group-specific overlays of individual connectivity distributions for (**A**) V2/3-TVA and (**B**) FFA-TVA connections, in red and green color respectively. The Arcuate Fasciculus (AF), Inferior Longitudinal Fasciculus (ILF), Inferior Fronto-Occipital Fasciculus (IFOF) were reconstructed with virtual *in vivo* interactive dissection (Catani et al. 2002) in a representative HC subject and depicted as anatomical landmarks (Catani and Thiebaut de Schotten 2008).

**Figure 3.**
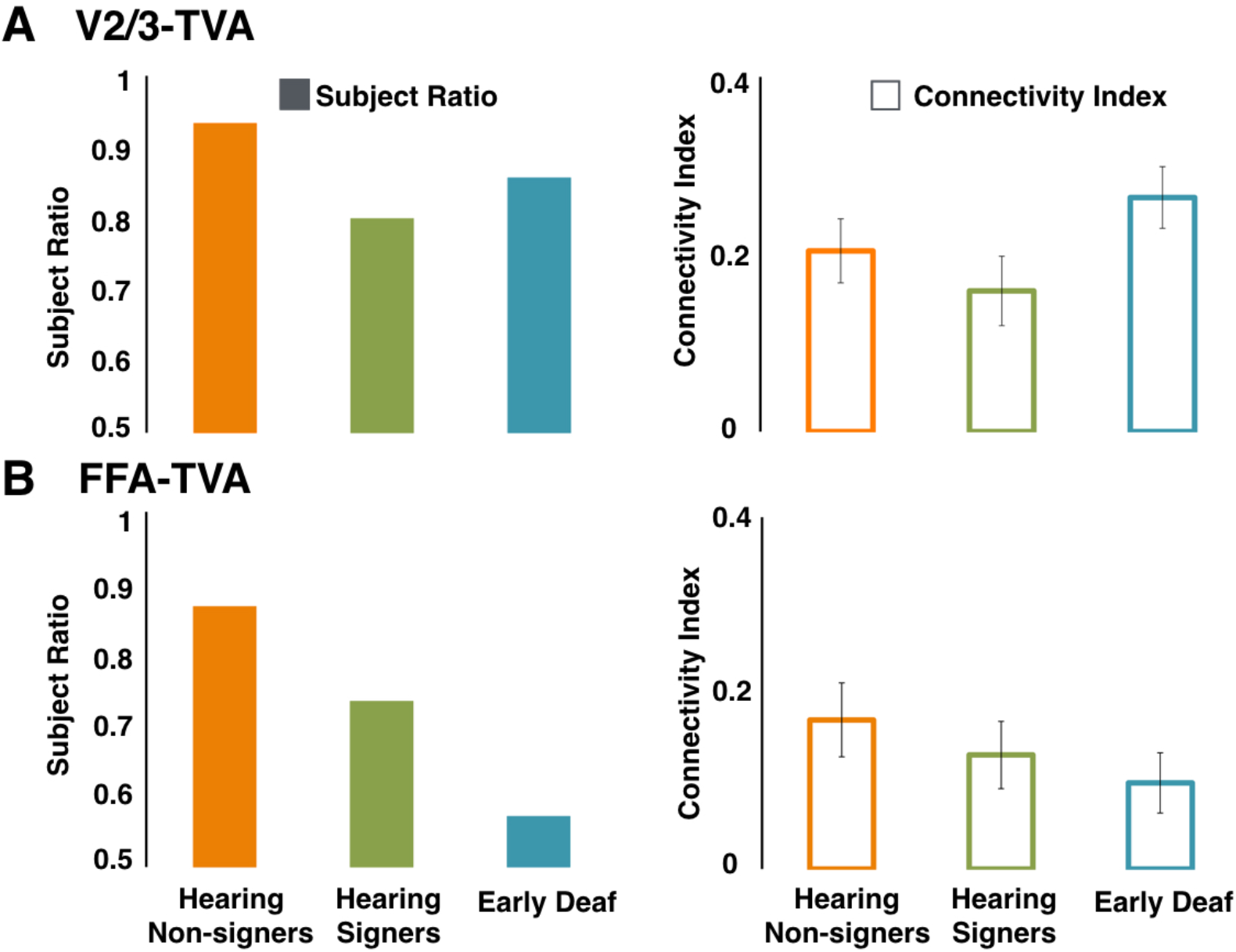
Quantitative analyses of macrostructural aspects of white matter pathways between (**A**) V2/3-TVA and (**B**) FFA-TVA areas. The plain bar-graphs depict the number of subjects showing a thresholded (> 10 streamlines) connection normalized by the number of all participants in each group. In each group, individual connectivity indices were calculated for each connection as the number of connected voxels divided by the overall number of connected voxels per participants and by the number of voxels of seed and target regions; the empty bar-graphs report group-averaged connectivity indices multiplied by 100 for presentation purposes. Error bars show SEM values. None of the pathways showed significant differences between groups (HC, orange; HC-SL, green; ED, cyan).

**Table 2.** (**da**) Macrostructural and (**b**) microstrνctural measures for the V2/3-TVA and FFA-TVA connections in the three experimental groups.

| (a) | V2/3 - TVA Pathway |  | FFA - TVA Pathway |  |
| --- | --- | --- | --- | --- |
| Group | Subject Ratio | Connectivity Index | Subject Ratio | Connectivity Index |
| Hearing Non-Signers | 0.93 | 0.203 (0.036) | 0.86 | 0.169 (0.041) |
| Hearing Signers | 0.80 | 0.158 (0.039) | 0.73 | 0.129 (0.038) |
| Early Deaf | 0.86+ | 0.263 (0.040)+ | 0.57+ | 0.097 (0.021)+ |

| (b) | V2/3 - TVA Pathway |  |  | FFA - TVA Pathway |  |  |
| --- | --- | --- | --- | --- | --- | --- |
| Group | FA | RD (x 10<sup>-3</sup>) | AD (x 10<sup>-3</sup>) | FA | RD (x 10<sup>-3</sup>) | AD (x 10<sup>-3</sup>) |
| Hearing Non-Signers | 0.371(0.007) | 0.56(0.01) | 0.99(0.01) | 0.374(0.007) | 0.55(0.01) | 0.99(0.01) |
| Hearing Signers | 0.390(0.007) | 0.54(0.01) | 0.99(0.01) | 0.386(0.007) | 0.53(0.01) | 0.98(0.02) |
| Early Deaf | 0.341(0.009)* | 0.62(0.02)* | 1.01(0.02) | 0.348(0.01)* | 0.61(0.01)* | 1.03(0.02) |

Standard error of the mean is reported in brackets. V2/3, extrastriate areas V2-V3; FFA, Face Fusiform Areas; TVA, Temporal Voice Area; FA, Fractional Anisotropy; RD, Radial Diffusivity; AD, Axial Diffusivity. (\*) significant group difference, $p < 0.025$ ; (+) Trend for significant difference in V2/3-TVA vs. FFA-TVA in Early Deaf, $p < 0.01$ . Significance is reported after Holm-Bonferroni correction as described in Methods.

In terms of connectivity strength indices, there were no significant differences between V2/3-TVA and FFA-TVA connections across groups (Wilcoxon Test, Z = 1.67, *p* = 0.095); this observation was also confirmed within the HC and HC-SL groups while the ED group showed a weak trend for higher connectivity strength in V2/3-TVA compared to FFA-TVA (Z = 1.98, *p* = 0.096, Table 2). Between the three groups, connectivity indices of V2/3-TVA and FFA-TVA were assessed with Kruskal-Wallis Tests that reported no significant group differences (Z = 3.62, *p* = 0.162 and Z = 2.55, *p* = 0.280, respectively, Table 2a).

### White matter microstructure of V2/3-TVA and FFA-TVA connections

Figure 4 shows the results of the microstructural measures comparison across groups and connections. Across the two white matter connections, the statistical analysis revealed a main effect of group (F_(2,22)_ = 4.72, *p* = 0.020, *η^2^* = 0.3) reflecting an overall reduction of FA in the ED group compared to HC-SL (*p* = 0.006) or HC (*p* = 0.055). Furthermore, differences in FA indices were associated with a significant increase of RD values (F_(2,21)_ = 6.224, *p* = 0.008, *η^2^* = 0.372) across connections in the ED group compared to both HC (*p* = 0.004) and HC-SL (*p* = 0.002) participants. No significant group differences were detected between the two hearing groups either for FA (FFA-TVA, *t* = −1.231, *p* = 0.688; V2/3-TVA, *t* = 2.073, *p* = 0.139) or RD (FFA-TVA, *t* = −0.713, *p* > 0.5; V2/3-TVA, *t* = 0.585, *p* = >0.5) values. No significant main effects of Connection (FA: F_(1,22)_ = 1.415, *p* = 0.247; RD: F_(1,21)_ = 0.001, *p* = 0.992) or Group-by-Connection interactions (FA: F_(1,22)_ = 0.198, *p* = 0.822; RD: F_(1,21)_ = 0.546, *p* = 0.587) were observed for mean FA and mean RD indices. Finally, we detected neither any main effects of Group (F_(2,21)_ = 1.917, *p* = 0.172) or Connection (F_(2,21)_ = 2.617, *p* = 0.121) nor a Group-by-Connection interaction (F_(2,21)_ = 0.778, *p* = 0.472) for AD values.

**Figure 4.**
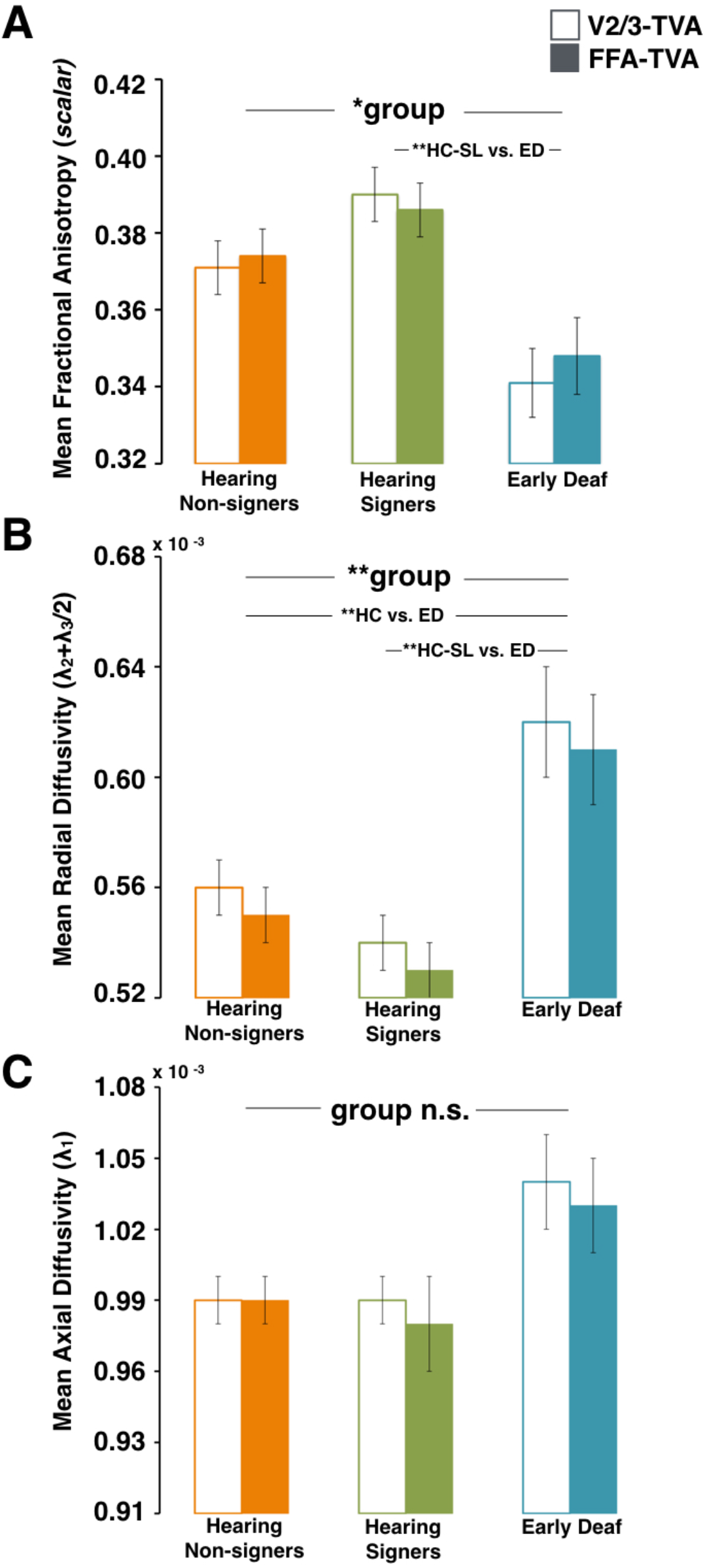
Quantitative analyses of microstructural aspects of V2/3-TVA (empty bars) and FFA-TVA (plain bars) connections. Significant differences were found for (**A**) fractional anisotropy and (**B**) radial diffusivity but not for (**C**) axial diffusivity in Early Deaf group relative to the Hearing groups. (**), *P* < 0.01; (*), *P* < 0.05. Radial and axial diffusivity values have been multiplied by 10^−3^ for display purposes. Error bars depict SEM values. Group-color scheme is the same as Fig. 3.

### Correlation between structural connectivity, functional TVA responses and behavioral measures

In the ED group, individual measures of face-selective activity in the right TVA (Benetti et al. 2017) showed no significant correlation either with FFA-TVA_(FA)_ indices or with FFA-TVA_(RD)_ values (R_(FA)_ = −0.658, *p* = 0.152; R_(RD)_ = 0.572, *p* = 0.207). No significant correlations were observed between diffusivity indices of the V2/3-TVA tract and face selective responses in the right TVA of ED individuals. While no significant correlations were revealed between microstructural indices and behavioral performances on face recognition across groups, in deaf individuals we detected trends for a negative correlation between individual face-recognition scores and FFA-TVA_(FA)_ (R = −0.732, *p* = 0.080) as well as a negative correlation with FFA-TVA_(RD)_ and V2/3-TVA_(RD)_ values (R = 0.709 *p* = 0.072 and R= 0.543, *p* = 0.060 respectively). When the HC-SL and ED groups were merged to assess the relationship between connectivity indices and LIS exposure no significant correlations were detected; similarly, there was no evidence of significant relationships between LIS exposure and either face-selective responses or face-recognition scores.

## Discussion

In this study, we implemented a multimodal imaging approach to examine whether the selective response to faces in the ‘deaf’ TVA and increased feed-forward effective connectivity between early visual extrastriate regions and the reorganized TVA observed in our previous work (Benetti et al., 2017) are associated with changes in white-matter connections within the face-voice brain network in the same deaf individuals. In summary, we were able to reconstruct both the V2/3-TVA and FFA-TVA connections in each experimental group. There were no significant differences between groups and between tracts for macrostructural measures of connectivity strength, although the ED group showed a weak trend towards higher connectivity strength in the V2/3-TVA compared to the FFA-TVA connection. At the microstructural level, we observed an overall reduction of mean FA values in ED participants compared to the two hearing groups, which was paralleled by a robust increase of mean RD indices across connections but not by differences in mean AD values. Finally, the correlation analyses revealed only a trend for significant relationship between FA/RD indices of both FFA-TVA and V2/3-TVA connections and face-recognition scores in deaf individuals, while no association between FA/RD indices and functional responses in right TVA was observed.

### Macrostructural connectivity

Our findings provide evidence for a occipito-temporal V2/3-TVA pathway, which was detected with high probability (≥80%) in each experimental group. Our reconstruction of V2/3-TVA pathway seems to overlap, at least in its medial and temporal components, with the dorsal bundles of the inferior longitudinal and inferior fronto-occipital fasciculi. These pathways connect occipital with temporal and frontal lobes, and are thought to be involved in face processing and visual perception respectively (Catani and Thiebaut de Schotten 2008). Whilst initial evidence for direct connections between visual and auditory regions have been reported (Beer et al. 2011), to the best of our knowledge, the present study is the first describing such connections between extrastriate visual and voice sensitive areas in humans. Previous animal studies have mostly reported direct reciprocal projections between visual areas V1/V2 and parabelt/caudal auditory regions in hearing monkeys and marmosets (Falchier et al. 2002, 2010; Rockland and Ojima 2003; Cappe and Barone 2005), which have been related to early audiovisual interactions (Cappe et al. 2009; Falchier et al. 2010). Indeed, functional dependencies between sensory systems exist between the earliest stages of sensory processing in both human (Ghazanfar and Schroeder 2006; Kayser et al. 2008; Schroeder and Lakatos 2009; Murray et al. 2016) and nonhuman primates (Lakatos et al., 2007; Schroeder and Lakatos, 2009). Early integration between facial and vocal information during the processing of speech may be supported by these direct connections between occipital and temporal regions (van Wassenhove et al. 2005), potentially relating to robust multisensory phenomenon like the McGurk effect (Saint-Amour et al. 2007). It has recently been suggested that the enhanced response to speech observed in the occipital regions of early blind may build on these intrinsic occipitotemporal connections (Van Ackeren et al. 2018). Likewise, it is therefore possible that the enhanced response to faces observed in the temporal cortex of congenitally deaf partially build on those connections as disclosed in our study. The high probability of detecting direct structural connectivity between V2/3 and TVA in the right hemisphere fits well with our previous observation of significant effective connectivity between these regions during face processing in deaf individuals, supporting our hypothesis that this heteromodal connection might be primarily involved in visual cross-modal reorganization of the deaf TVA (Benetti et al. 2017).

The detection of a direct link between FFA and TVA in each group is also in line with previous findings, which reported both functional (von Kriegstein et al. 2005) and structural (Blank et al. 2011) connectivity between right FFA and voice-sensitive regions in the mid/anterior-STG of hearing individuals (Fig 2B). It has been suggested that direct reciprocal connections between FFA and TVA play a key role in exchanging predictive face-voice features that are relevant for identity recognition and that can be used to constrain ongoing interpretations of unisensory ambiguous inputs (Blank et al. 2011). Although no significant differences were observed between groups, the reconstruction of FFA-TVA connections was possible for fewer ED than hearing participants (57% *vs*. 86%). Since no differences were reported in data quality (see Methods and Limitations sections), one tentative interpretation would relate this observation in the deaf participants to a potential differential experience-dependent refining of this anatomical pathway due to differences in the contribution of V2/3-TVA and FFA-TVA connections to face-voice processing. For instance, FFA-TVA refining might be more sensitive to an early lack of coincidence between face and voice inputs during identity encoding and maintenance in deaf individuals (von Kriegstein et al. 2005; Schall et al. 2013).

The examination of macrostructural connectivity strength did not reveal significant group differences for either pathway. Previous investigations of cortico-cortical connectivity in congenitally or early deaf cats (ototoxically deafened < 50 days of life) have most consistently revealed that area A1 (Barone et al. 2013; Chabot et al. 2015), area A2 (Butler et al. 2017b), the Dorsal Zone (Barone et al. 2013), the auditory field of the Anterior Ectosylvian Sulcus (Meredith et al. 2016) and the Posterior Auditory Field (Butler et al. 2017a) receive additional visual and somatosensory projections (as measured by injections of anatomical tracers) that are absent in hearing cats. These new projections, however, represent only a small fraction of the inputs to the auditory regions; moreover, their areal specificity and strength overall appear comparable to those observed for normal projections in hearing cats (Barone et al. 2013; Chabot et al. 2015; Meredith et al. 2016; Butler et al. 2017b). Accordingly, it has been suggested that the emergence of functional cross-modal plasticity might be, at least in part, supported by a combination of both preserved heteromodal structural connections and minor reorganization of cortico-cortical connectivity in early deafness (Barone et al. 2013; Butler et al. 2017b). This notion is also supported by the evidence, provided by animal models of blindness, that a near-normal pattern of connections is maintained in the occipital cortex in the absence, or following early lack, of visual inputs (Berman 1991; Karlen et al. 2006). In addition, a similar dichotomy between near-normal patterns of anatomical connectivity and substantial functional reorganization has also been reported in visually deprived cats (Mower et al. 1985) and early blind humans (Shimony et al. 2006). Our macrostructural observations, while consistent with the notion of preserved heteromodal structural connectivity, does not provide support to the hypothesis that additional visual projections sustain visual recruitment of the right TVA in congenital deaf humans. Alternatively, they might suggest that the development of such additional projections is not reflected by changes in macrostructural connectivity between visual and auditory regions typically involved in face and voice processing.

### Microstructural connectivity

In contrary to macrostructural connectivity, the examination of microstructural diffusivity measures revealed an overall pattern of alterations in the deaf relative to the hearing individuals. When comparing early deaf with controls we found on both pathways a significant reduction of FA that was associated with an increase of RD but no alterations of AD. To date, a handful of previous studies have examined microstructural diffusivity measures in deaf humans by applying tensor imaging techniques to cortico-cortical connections, either with whole brain or region of interest approaches. Only one study investigated diffusivity measures within long-range projections, but focused on thalamo-cortical auditory connectivity (Lyness et al. 2014). Our observation of reduced FA is in line with the four studies that examined whole brain connectivity and consistently reported reductions of FA in the superior temporal cortex of deaf individuals (Kim et al. 2009; Li et al. 2012; Miao et al. 2013; Hribar et al. 2014). Our findings seems also consistent with a recent study (Karns et al. 2017) that focused on regions of interest within the superior temporal cortex (i.e. Heschl’s gyrus and anterior/posterior STG) and reported overall FA reductions associated with increased RD and unchanged AD measures.

The relationship between changes in diffusivity indices and alterations in the underlying biological microstructure is not straight-forward (Beherens et al., 2013; Wheeler-Kingshott and Cercignani 2009). Fractional anisotropy measures reflect the directional hindering of diffusivity of water molecules in white matter bundles. Reductions of this index have been related to altered integrity of the microstructural aspects (e.g. axonal packing, axonal thickness and myelination) of white matter bundles (Alexander et al. 2007; Assaf and Pasternak 2008). Radial and axial diffusivity indices, that have been related to reduced myelination and microtubule deficits in studies of white matter pathologies (Alexander et al. 2007; Behrens et al. 2013), provide additional information on diffusivity along the different fiber axis and can aid the interpretation of FA reductions. Alterations in axonal myelination, which are reflected by FA reductions, can be associated with increased RD and reduced or unchanged AD (Song et al. 2002). However, when speculating about radial and axial diffusivity changes, a univocal association with specific biological changes can prove difficult in the absence of histological examinations and should be cautiously considered (Wheeler-Kingshott and Cercignani 2009). We therefore propose two plausible interpretations for the observed between-groups differences in diffusivity configuration. In deaf humans, the observed configuration might result from either atypical myelination or subtle axonal atrophy as a consequence of lack of auditory stimulation and processing during brain and deafness development (Karns et al. 2017). Indeed, a number of studies consistently reported reductions of white matter volume in auditory and their neighboring temporal regions in early deafness despite preserved gray matter volume (Emmorey et al. 2003; Shibata 2007; Smith et al. 2011; Hribar et al. 2014). Similarly, reductions of white matter volume have also been reported in visual regions of individuals with early visual deprivation (Noppeney et al. 2005; Ptito et al. 2008). Although the precise cellular mechanisms supporting white matter changes are currently under debate, there is growing evidence that experience-dependent activity can induce myelination changes in the adult and developing brain (for a recent review, Sampaio-Baptista and Johansen-Berg 2017). In particular, Etxeberria and colleagues reported shortened myelin sheathes and reduce axonal impulse conduction in visually deprived mice during their development (Etxeberria et al. 2016). In our sample, atypical development might have selectively affected the feedback projections connecting the temporal voice area to high-order extrastriate and fusiform visual areas due to the lack of vocal input within the face-voice network. Feedback projections appear late in the cortical development and have longer developmental times, which makes them more likely to be affected by sensory experience (Kral et al. 2017). Previous research on animal models of deafness, for example, reported reduced activity (Kral et al. 2000) along with a subtle reduction of thickness (Berger et al. 2017), extending from primary to secondary auditory regions, mostly in the deep infragranular layers. These layers are the substrate of top-down interactions in the cortical columnar architecture (Barone et al. 2000; Grossberg 2007), therefore, it is likely that such functional and morphological changes reflect alterations of the feedback projections arising from infragranular layers.

While the lack of auditory inputs and voice-dependent activity in the deaf sample can provide a plausible explanation for our observations at the microstructural level, a second possibility is that increased RD and reduced FA indices reflect a greater white matter complexity tissue driven by an increased number of crossing fibers, especially at the boundaries with gray matter (Wheeler-Kingshott et al. 2009), that could be associated with cross-modal recruitment of the auditory regions (Karns et al. 2017). In fact, the comparative study of diffusion properties and tissue histology in the human brain suggests that decreases in FA might be observed even after learning or training and reflect the maturation of secondary crossing fiber populations (for a review see Zatorre et al. 2012). According to this notion, in our deaf sample greater crossing fibers might be associated with increased cross-modal recruitment and functional connectivity between occipital and temporal regions.

These two interpretations, atypical development and greater tissue complexity due to cross-modal plasticity, are not mutually exclusive. Our observation of trends for correlation between microstructural indices (RD = positive and FA = negative) and face recognition scores for both occipito- and fusiform-temporal connections might support the hypothesis of greater tissue complexity induced by (compensative) cross-modal plasticity. However, a direct causal relationship between connectivity indices and behavioral performances cannot be easily inferred. For instance, a supra-normal ability in visual face recognition might also be supported by local changes in synaptic strength and efficiency of existing visual projections (Rauschecker 1995; Bavelier and Neville 2002) in concomitance to, and despite of, an atypical development of feedback auditory projections associated with early lack of acoustic input from auditory to visual regions. Therefore, the two hypotheses remain overall untested and could be further addressed in animal models of deafness.

### Limitations

As for all studies involving diffusion MRI data, a number of potential caveats need to be taken into consideration when interpreting and reconciling them with functional MRI data.

In the present study we found no structural connectivity changes related to the increased functional connectivity previously reported within the reorganized face-voice system of deaf individuals (Benetti et al. 2017). Such discrepancies between structural and functional results have been previously reported in multimodal imaging studies (Benetti et al. 2015; Bauer et al. 2017) and may be due to several reasons. Unlike diffusion-based structural connectivity that characterizes physical pathways between regions, functional connectivity quantifies the statistical dependencies of structures (cross-correlation) and depends on estimated regional activity expressed as activity-dependent time-series. Whilst increased functional connectivity between two regions may be indicative of an underlying structural pathway, its extent may depend also on indirect structural connections between the same regions. For instance, from our previous observations, we cannot exclude that the influence exerted by V2/3 activity on face-selectivity in TVA was mediated by other regions not included in our dynamical causal modeling (Benetti et al. 2017).

There are also several factors that can potentially affect the results of diffusion-based tractography studies. Poor quality of data often results in a failure to reconstruct paths between cortical regions, rather than introducing systematic error (Behrens et al. 2003). In this study, we implemented an *ad-hoc* dMRI sequence with the aim to improve the signal-to-noise ratio and avoid implausible tracking. Then, the acquisition of dMRI/fMRI data was carried out in a pseudorandomized manner between groups in order to avoid that potential fluctuations of data quality would impact one single group. Further, individual dMRI images were visually inspected for artifacts and the signal-to-noise ratio checked (see methods), suggesting comparable degrees of data quality across groups. A second potential issue relates to the complexity and geometry of the underlying biological structure (Johansen-Berg et al. 2009), whereby tracking paths with high curvatures and potential crossing-fiber or close to grey matter can result in limited connectivity findings. We, therefore, used probabilistic multi-fiber tracking, which provides more sensitivity to crossing fibers (Behrens et al. 2007). In addition, for our functionally defined seed/target regions, we tracked only from proximal white matter voxels, which has proved successful in previous dissection of structural paths between face-voice areas (Blank et al. 2011). The physical distance of the seed to the target regions may also impact on the ease of path tracking since connection between closer regions may be associated with greater probability of tracking (Tomassini et al. 2007). However, we did not observed differences between tracts across groups, suggesting that cortical target proximity was unlikely to have introduced systematic distortions in this investigation. It is also important to bear in mind a number of potential limitations that applies to diffusion studies and may relate to specific experimental aspects of our study, such as the spatial resolution and diffusion metrics chosen for the diffusion analysis. For instance, structural changes in deaf individuals might occur at a higher spatial resolution or be reflected by tissue properties that were not measurable by the diffusion metrics computed in this study, or the combination of the two. Nevertheless, our findings seems to be consistent with alterations also reported by previous human deaf studies (Kim et al. 2009; Hribar et al. 2014; Karns et al. 2017) and anatomical tracer studies in deaf animals (Barone et al. 2013; Meredith et al. 2016). In the following section, therefore, we will attempt a cautious reconciliation of our structural and functional observations by relating them to existing evidence from human and animal studies.

### Interpreting functional and structural observations in the context of functionally selective neuroplasticity

In the present study, we examined the structural connectivity within a face-voice system, which presents local and large-scale functionally selective crossmodal reorganization in congenitally and early deaf humans (Benetti et al. 2017). We report only changes, in early deaf relative to normally hearing subjects, in the microstructural connectivity between extrastriate and fusiform face-responsive regions to the temporal voice area in the right hemisphere. Overall our observations support the notion that local and large-scale functional reorganization is not directly reflected in the underlying pattern of white matter connectivity (Bauer et al. 2017; Butler et al. 2017b; Karns et al. 2017; Kral et al. 2017). We propose that a mixed contribution of genetic and both experience-dependent loss and compensatory mechanisms induces differential changes within the structural architecture of the reorganized face-voice system, yielding the anatomo-functional configuration associated with face-selective cross-modal reorganization of the ‘deaf’ temporal voice area (Benetti et al., 2017). In other words, we presume that the existence of innate anatomical networks between sensory and cognitive systems constrains the expression of crossmodal plasticity by inducing a reorganization process (e.g. enhanced visual responses in the temporal cortex of the deaf) that follows functional principles derived from such intrinsic anatomical connections. In particular, the existence of anatomical pathways linking visual and auditory regions implicated in face and voice processing (see Fig. 1 and 2) would constraint the functional remapping of faces in the deprived temporal voice area in contrast to other visual stimuli (e.g. houses; see Benetti et al. 2017).

Indeed, the early anatomical development of white matter bundles is mostly dependent on genetic control and molecular guiding factors (Innocenti et al. 1988; Innocenti and Price 2005), which might prescribe the general trajectory and response to environmental and experience related factors (Johansen-Berg and Behrens 2013). However, experience-dependent activity can modulate early selective myelination of fiber axons; thus, it plays an important role in the fine-tuning of cortico-cortical connections (Price et al. 2006), more relevantly of connections supporting integration of multisensory information (Kral et al. 2017). Relevantly, it has been proposed that the emergence of category-selective responses (such as face-selectivity) within a specialized cortical network is enhanced by experienced-dependent feedback projections (Reid 2012) but it is guided by pre-existing patterns of anatomical connectivity, which provide a scaffolding for the subsequent developmental refinement of responses (Kanwisher 2010; Hannagan et al. 2015; Deen et al. 2017). Our study seems to suggest that the lack of early voice-evoked neural activity may impact only on the microstructural anatomical connectivity within the functionally reorganized face-voice system. The maintenance of macrostructural connectivity, therefore, might provide the structural substrate for feeding of early facial-related visual information into the deafened temporal voice area; accordingly, the evidence of a direct connection between occipital and temporal regions is consistent with increased functional coupling and significant effective connectivity between these regions (Benetti et al. 2017). Preservation of occipito-temporal structural pathways might, therefore, allow ‘deafened’ temporal areas to participate preferentially in some specific visual functions (Land et al. 2016). The notion that the stabilization and maintenance of cortico-cortical connectivity might constrain the expression of cross-modal plasticity following sensory deprivation is not confined to deafness models, but has long been suggested by observations reported in blind humans (Noppeney 2007; Mahon and Caramazza 2011; Collignon et al. 2011). Recent investigations, which focused specifically on effective functional connectivity in blind individuals, also showed that auditory colonization of visual regions is better explained by direct cortico-cortical connections with primary auditory cortex in congenitally (Klinge et al. 2010) but not late blind (Collignon et al. 2013) individuals, further supporting the hypothesis that experiential factors associated with sensory experience interact with predetermined genetic programs for the development and fine tuning of long-range cortico-cortical connectivity.

An alternative hypothesis would be that cortical reorganization might depend on rearrangement of subcortical inputs. For example, there is recent evidence in favor of conversion from auditory to somatosensory thalamic input (Allman et al. 2009), as well as increased somatosensory projections to auditory regions (Zeng et al. 2012) in deaf animals. In our previous functional investigation, no face-selective reorganization of thalamo-cortical coupling was found (Benetti et al. 2017). This suggested, that cross-modal selectivity to high-order visual stimuli, such as human faces, is unlikely to be specifically driven by rearrangement of thalamo-cortical visual inputs. Consistently, a recent tractography study failed to report alterations in the thalamo-cortical connections of deaf individuals (Lyness et al. 2014), suggesting that profound deafness does not extensively impact on subcortical connections (see Klinge et al. 2010 for a similar proposal in blindness).

In the context of increased cortico-cortical functional connectivity and unchanged, or even slightly reduced, structural connectivity a more plausible and parsimonious interpretation could be favored. Rather than originating from long-range anatomical reorganization within the face-voice system, functionally selective recruitment of auditory regions might result from changes in synaptic distribution, strength and efficiency of existing projections (Rauschecker 1995; Bavelier and Neville 2002; Collignon et al. 2009). In fact, Hebbian mechanisms of plasticity (e.g. long term potentiation/depression) are synaptically controlled and thought to primarily induce changes at the level of dendritic pre- and post-synaptic sites (Siegelbaum and Kandel 1991; Magee and Johnston 1997); accordingly, the effective modulation that one brain area can exert over another is thought to be mostly defined by the synaptic counts and strengths (Friston 2011). It has been proposed that such synaptic reorganization might affect existing projections that normally provide sub-threshold cross-modal inputs to the auditory regions (Butler et al. 2017b; Kral et al. 2017). Indeed, in addition to the typical selective response to human voice, we observed a subtle, yet not significant face selective response in the voice area of hearing participants (Benetti et al. 2017), pointing to the contribution of heteromodal projections in supporting multisensory integration. This hypothesis has received support in recent studies relying on animal models of deafness. For instance, increases in the spine density of neurons have been reported in early and high-order auditory regions of deaf cats (Clemo et al. 2016, 2017). Further, the study of deaf cats with cochlear implants has shown that visually-responsive neurons are spatially scattered between auditory-responsive neurons in the auditory regions, suggesting increased synaptic development of existing projections that were strengthen (Land et al. 2016). It should be noted that, although we do not report evidence of significant relationships between LIS exposure and connectivity measures or differences between the two hearing groups, specific aspects of sign language processing (such as increased reliance on dynamic visual face and lip cues) might also have modulated the effects of auditory deprivation on the organization of the face/voice system in the deaf and hearing signers included in this study.

In conclusion, our study is the first attempt to integrate anatomo-functional observations within a neural system showing task-specific, large-scale cross-modal reorganization in humans with sensory deprivation. Our main findings are overall in line with evidence from animal models (Barone et al. 2013; Clemo et al. 2016; Meredith et al. 2016; Butler et al. 2017b) suggesting a maintenance of white matter connections between visual and auditory cortices following deafness and supporting the idea that innate anatomical networks between sensory/cognitive systems constrain the expression of crossmodal plasticity. We further propose that genetic and experiential factors might differentially impact on the development and maintenance of the structural network within the face-voice processing system. In general, we suggest that further efforts are needed to better understand how cortical structure and function, white matter connectivity and behavior are linked in cross-modal plasticity. To this aim, it is crucial to combine multimodal imaging information within specific sensory and cognitive systems showing functional selective reorganization, and to integrate it with individual measures of behavioral performance.

## Acknowledgments

We thank all the deaf and hearing people who participated in this research for their collaboration. We equally thank Valentina Foa, Joshua Zonca and Francesca Baruffaldi for their support with subject recruitment and testing, as well as throughout completion of this study. O.C. is a research associate at the Fond National de la Recherche Scientifique of Belgium (FRS-FNRS).

## Ethical Approval

The study was approved by the Committee for Research Ethics of the University of Trento; all participants gave informed consent in agreement with the ethical principles for medical research involving human subject (Declaration of Helsinki, World Medical Association) and the Italian Law on individual privacy (D.l. 196/2003).

## References

Abdi H (2010) Holm’s Sequential Bonferroni Procedure. In: Encyclopedia of Research Design. pp 1–8

Alexander AL, Lee JE, Lazar M, Field AS (2007) Diffusion tensor imaging of the brain. Neurotherapeutics 4:316–29. doi: 10.1016/j.nurt.2007.05.011

Allman BL, Keniston LP, Meredith MA (2009) Adult deafness induces somatosensory conversion of ferret auditory cortex. Proc Natl Acad Sci 106:5925–5930. doi: 10.1073/pnas.0809483106

Assaf Y, Pasternak O (2008) Diffusion tensor imaging (DTI)-based white matter mapping in brain research: A review. J. Mol. Neurosci. 34:51–61

Auer ET, Bernstein LE, Sungkarat W, Singh M (2007) Vibrotactile activation of the auditory cortices in deaf versus hearing adults. Neuroreport 18:645–648

Barone P, Batardiere A, Knoblauch K, Kennedy H (2000) Laminar distribution of neurons in extrastriate areas projecting to visual areas V1 and V4 correlates with the hierarchical rank and indicates the operation of a distance rule. J Neurosci 20:3263–3281. doi: 10.1523/jneurosci.0414-12.2012

Barone P, Lacassagne L, Kral A (2013) Reorganization of the Connectivity of Cortical Field DZ in Congenitally Deaf Cat. PLoS One 8: doi: 10.1371/journal.pone.0060093

Basser PJ, Pierpaoli C (2011) Microstructural and physiological features of tissues elucidated by quantitative-diffusion-tensor MRI. J Magn Reson 213:560–570. doi: 10.1016/j.jmr.2011.09.022

Bauer CM, Hirsch G V., Zajac L, et al (2017) Multimodal MR-imaging reveals large-scale structural and functional connectivity changes in profound early blindness. PLoS One 12: doi: 10.1371/journal.pone.0173064

Bavelier D, Neville HJ (2002) Cross-modal plasticity: where and how? Nat Rev Neurosci 3:443–452

Beer AL, Plank T, Greenlee MW (2011) Diffusion tensor imaging shows white matter tracts between human auditory and visual cortex. Exp brain Res 213:299–308

Behrens TEJ, Berg HJ, Jbabdi S, et al (2007) Probabilistic diffusion tractography with multiple fibre orientations: What can we gain? Neuroimage 34:144–155. doi: 10.1016/j.neuroimage.2006.09.018

Behrens TEJ, Sotiropoulos SN, Jbabdi S (2013) MR Diffusion Tractography. In: Diffusion MRI: From Quantitative Measurement to In vivo Neuroanatomy: Second Edition. pp 429–451

Behrens TEJ, Woolrich MW, Jenkinson M, et al (2003) Characterization and Propagation of Uncertainty in Diffusion-Weighted MR Imaging. Magn Reson Med 50:1077–1088. doi: 10.1002/mrm.10609

Belin P, Zatorre RJ, Lafaille P, et al (2000) Voice-selective areas in human auditory cortex. Nature 403:309–312

Benetti S, Pettersson-Yeo W, Allen P, et al (2015) Auditory verbal hallucinations and brain dysconnectivity in the perisylvian language network: A multimodal investigation. Schizophr Bull 41:192–200

Benetti S, van Ackeren MJ, Rabini G, et al (2017) Functional selectivity for face processing in the temporal voice area of early deaf individuals. Proc Natl Acad Sci 114:E6437–E6446. doi: 10.1073/pnas.1618287114

Berger C, Kühne D, Scheper V, Kral A (2017) Congenital deafness affects deep layers in primary and secondary auditory cortex. J Comp Neurol 525:3110–3125. doi: 10.1002/cne.24267

Berman NEJ (1991) Alterations of visual cortical connections in cats following early removal of retinal input. Dev Brain Res 63:163–180. doi: 10.1016/0165–3806(91)90076-U

Blank H, Anwander A, von Kriegstein K (2011) Direct structural connections between voice- and face-recognition areas. J Neurosci 31:12906–12915

Blank H, Kiebel SJ, von Kriegstein K (2015) How the human brain exchanges information across sensory modalities to recognize other people. Hum Brain Mapp 36:324–339. doi: 10.1002/hbm.22631

Bola Ł, Zimmermann M, Mostowski P, et al (2017) Task-specific reorganization of the auditory cortex in deaf humans. Proc Natl Acad Sci Early Edit: 1–10. doi: 10.1073/pnas.1609000114

Butler BE, Chabot N, Kral A, Lomber SG (2017a) Origins of thalamic and cortical projections to the posterior auditory field in congenitally deaf cats. Hear Res 343:118–127. doi: 10.1016/j.heares.2016.06.003

Butler BE, de la Rua A, Ward-Able T, Lomber SG (2017b) Cortical and thalamic connectivity to the second auditory cortex of the cat is resilient to the onset of deafness. Brain Struct. Funct. 1–17

Cappe C, Barone P (2005) Heteromodal connections supporting multisensory integration at low levels of cortical processing in the monkey. Eur J Neurosci 22:2886–2902. doi: 10.1111/j.1460–9568.2005.04462.x

Cappe C, Rouiller EM, Barone P (2009) Multisensory anatomical pathways. Hear Res 258:28–36. doi: 10.1016/j.heares.2009.04.017

Catani M, Howard RJ, Pajevic S, Jones DK (2002) Virtual in vivo interactive dissection of white matter fasciculi in the human brain. Elsevier B.V., Section of Old Age Psychiatry, Institute of Psychiatry, De Crespigny Park, London SE5 8AF, United Kingdom.

Catani M, Thiebaut de Schotten M (2008) A diffusion tensor imaging tractography atlas for virtual in vivo dissections. Cortex

Chabot N, Butler BE, Lomber SG (2015) Differential Modification of Cortical and Thalamic Projections to Cat Primary Auditory Cortex Following Early- and Late-Onset Deafness. J Comp Neurol 523:2297–320. doi: 10.1002/cne.23790

Choi S, Cunningham DT, Aguila F, et al (2011) DTI at 7 and 3 T: Systematic comparison of SNR and its influence on quantitative metrics. Magn Reson Imaging 29:739–751. doi: 10.1016/j.mri.2011.02.009

Clemo HR, Lomber SG, Meredith MA (2016) Synaptic Basis for Cross-modal Plasticity: Enhanced Supragranular Dendritic Spine Density in Anterior Ectosylvian Auditory Cortex of the Early Deaf Cat. Cereb Cortex 26:1365–1376. doi: 10.1093/cercor/bhu225

Clemo HR, Lomber SG, Meredith MA (2017) Synaptic distribution and plasticity in primary auditory cortex (A1) exhibits laminar and cell-specific changes in the deaf. Hear. Res.

Collignon O, Dormal G, Albouy G, et al (2013) Impact of blindness onset on the functional organization and the connectivity of the occipital cortex. Brain 136:2769–2783. doi: 10.1093/brain/awt176

Collignon O, Vandewalle G, Voss P, et al (2011) Functional specialization for auditory-spatial processing in the occipital cortex of congenitally blind humans. Proc Natl Acad Sci U S A 108:4435–4440

Collignon O, Voss P, Lassonde M, Lepore F (2009) Cross-modal plasticity for the spatial processing of sounds in visually deprived subjects. Exp brain Res Exp Hirnforsch Expérimentation cérébrale 192:343–358

Deen B, Richardson H, Dilks DD, et al (2017) Organization of high-level visual cortex in human infants. Nat Commun 8: doi: 10.1038/ncomms13995

Dormal G, Collignon O (2011) Functional selectivity in sensory-deprived cortices. J Neurophysiol

Doron KW, Funk CM, Glickstein M (2010) Fronto-cerebellar circuits and eye movement control: A diffusion imaging tractography study of human cortico-pontine projections. Brain Res 1307:63–71. doi: 10.1016/j.brainres.2009.10.029

Eickhoff SB, Jbabdi S, Caspers S, et al (2010) Anatomical and functional connectivity of cytoarchitectonic areas within the human parietal operculum. J Neurosci 30:6409–6421. doi: 10.1523/jneurosci.5664–09.2010

Emmorey K, Allen JS, Bruss J, et al (2003) A morphometric analysis of auditory brain regions in congenitally deaf adults. Proc Natl Acad Sci U S A 100:10049–10054

Etxeberria A, Hokanson KC, Dao DQ, et al (2016) Dynamic Modulation of Myelination in Response to Visual Stimuli Alters Optic Nerve Conduction Velocity. J Neurosci 36:6937–48. doi: 10.1523/JNEUROSCI.0908–16.2016

Falchier A, Clavagnier S, Barone P, Kennedy H (2002) Anatomical evidence of multimodal integration in primate striate cortex. J Neurosci 22:5749–5759

Falchier A, Schroeder CE, Hackett TA, et al (2010) Projection from visual areas V2 and prostriata to caudal auditory cortex in the monkey. Cereb Cortex 20:1529–1538. doi: 10.1093/cercor/bhp213

Fillard P, Descoteaux M, Goh A, et al (2011) Quantitative evaluation of 10 tractography algorithms on a realistic diffusion MR phantom. Neuroimage 56:220–234. doi: 10.1016/j.neuroimage.2011.01.032

Fine I, Finney EM, Boynton GM, Dobkins KR (2005) Comparing the effects of auditory deprivation and sign language within the auditory and visual cortex. J Cogn Neurosci 17:1621–1637. doi: 10.1162/089892905774597173

Finney EM, Fine I, Dobkins KR (2001) Visual stimuli activate auditory cortex in the deaf. Nat Neurosci 4:1171–1173

Friston KJ (2011) Functional and Effective Connectivity: A Review. Brain Connect 1:13–36. doi: 10.1089/brain.2011.0008

Ghazanfar A, Schroeder C (2006) Is neocortex essentially multisensory? Trends Cogn Sci 10:278–285. doi: 10.1016/j.tics.2006.04.008

Grossberg S (2007) Towards a unified theory of neocortex: laminar cortical circuits for vision and cognition. Prog. Brain Res. 165:79–104

Hannagan T, Amedi A, Cohen L, et al (2015) Origins of the specialization for letters and numbers in ventral occipitotemporal cortex. Trends Cogn Sci 19:374–382

Heimler B, Weisz N, Collignon O (2014) Revisiting the adaptive and maladaptive effects of crossmodal plasticity. Neuroscience 283:44–63

Holm S (1979) A simple sequential rejective multiple test procedure. Scand J Stat 6:65–70. doi: 10.2307/4615733

Hribar M, Suput D, Carvalho AA, et al (2014) Structural alterations of brain grey and white matter in early deaf adults. Hear Res 318:10

Innocenti GM, Berbel P, Clarke S (1988) Development of projections from auditory to visual areas in the cat. J Comp Neurol 272:242–259. doi: 10.1002/cne.902720207

Innocenti GM, Price DJ (2005) Exuberance in the development of cortical networks. Nat. Rev. Neurosci. 6:955–965

Javad F, Warren JD, Micallef C, et al (2014) Auditory tracts identified with combined fMRI and diffusion tractography. Neuroimage 84:562–574. doi: 10.1016/j.neuroimage.2013.09.007

Johansen-Berg H, Behrens TEJ (2013) Diffusion MRI: From Quantitative Measurement to In vivo Neuroanatomy: Second Edition

Johansen-Berg H, Rushworth MF, Berg HJ, Rushworth MFS (2009) Using Diffusion Imaging to Study Human Connectional Anatomy. Annu Rev Neurosci 32:75–94. doi: 10.1146/annurev.neuro.051508.135735

Kanwisher N (2010) Functional specificity in the human brain: a window into the functional architecture of the mind. PNAS 107:11163–70. doi: 10.1073/pnas.1005062107

Kanwisher N, McDermott J, Chun MM (1997) The fusiform face area: a module in human extrastriate cortex specialized for face perception. J Neurosci 17:4302–11. doi: 10.1098/Rstb.2006.1934

Karlen SJ, Kahn DM, Krubitzer L (2006) Early blindness results in abnormal corticocortical and thalamocortical connections. Neuroscience 142:843–858. doi: 10.1016/j.neuroscience.2006.06.055

Karns CM, Dow MW, Neville HJ (2012) Altered cross-modal processing in the primary auditory cortex of congenitally deaf adults: a visual-somatosensory fMRI study with a double-flash illusion. J Neurosci 32:9626–9638

Karns CM, Stevens C, Dow MW, et al (2017) Atypical white-matter microstructure in congenitally deaf adults: A region of interest and tractography study using diffusion-tensor imaging. Hear Res 343:72–82. doi: 10.1016/j.heares.2016.07.008

Kayser C, Petkov CI, Logothetis NK (2008) Visual Modulation of Neurons in Auditory Cortex. Cereb Cortex 18:1560–1574. doi: 10.1093/cercor/bhm187

Kim D-J, Park S-Y, Kim J, et al (2009) Alterations of white matter diffusion anisotropy in early deafness. Neuroreport 20:1032–1036

Klinge C, Eippert F, Röder B, Büchel C (2010) Corticocortical connections mediate primary visual cortex responses to auditory stimulation in the blind. J Neurosci 30:12798–805. doi: 10.1523/JNEUROSCI.2384–10.2010

Kral A, Hartmann R, Tillein J, et al (2000) Congenital auditory deprivation reduces synaptic activity within the auditory cortex in a layer-specific manner. Cereb Cortex 10:714–726. doi: 10.1093/cercor/10.7.714

Kral A, Yusuf PA, Land R (2017) Higher-order auditory areas in congenital deafness: Top-down interactions and corticocortical decoupling. Hear. Res. 343:50–63

Lakatos P, Chen C, Connell MNO, et al (2007) Article Neuronal Oscillations and Multisensory Interaction in Primary Auditory Cortex. 279–292. doi: 10.1016/j.neuron.2006.12.011

Land R, Baumhoff P, Tillein J, et al (2016) Cross-Modal Plasticity in Higher-Order Auditory Cortex of Congenitally Deaf Cats Does Not Limit Auditory Responsiveness to Cochlear Implants. J Neurosci 36:6175–6185. doi: 10.1523/JNEUROSCI.0046-16.2016

Lao Y, Kang Y, Collignon O, et al (2015) A study of brain white matter plasticity in early blinds using tract-based spatial statistics and tract statistical analysis. Neuroreport 26:1151–1154. doi: 10.1097/WNR.0000000000000488

Leemans A, Jones DK (2009) The B-matrix must be rotated when correcting for subject motion in DTI data. Magn Reson Med 61:1336–1349. doi: 10.1002/mrm.21890

Li Y, Ding G, Booth JR, et al (2012) Sensitive period for white-matter connectivity of superior temporal cortex in deaf people. Hum Brain Mapp 33:349–359. doi: 10.1002/hbm.21215

Lomber SG, Meredith MA, Kral A (2010) Cross-modal plasticity in specific auditory cortices underlies visual compensations in the deaf. Nat Neurosci 13:1421–1427

Lyness RC, Alvarez I, Sereno MI, MacSweeney M (2014) Microstructural differences in the thalamus and thalamic radiations in the congenitally deaf. Neuroimage 100:347–357. doi: 10.1016/j.neuroimage.2014.05.077

Magee JC, Johnston D (1997) A synaptically controlled, associative signal for Hebbian plasticity in hippocampal neurons. Science (80-) 275:209–213. doi: 10.1126/science.275.5297.209

Mahon BZ, Caramazza A (2011) What drives the organization of object knowledge in the brain? Trends Cogn. Sci. 15:97–103

Makuuchi M, Bahlmann J, Anwander A, Friederici AD (2009) Segregating the core computational faculty of human language from working memory. Proc Natl Acad Sci 106:8362–8367. doi: 10.1073/pnas.0810928106

Meredith MA, Clemo HR, Corley SB, et al (2016) Cortical and thalamic connectivity of the auditory anterior ectosylvian cortex of early-deaf cats: Implications for neural mechanisms of crossmodal plasticity. Hear Res 333:25–36. doi: 10.1016/j.heares.2015.12.007

Miao W, Li J, Tang M, et al (2013) Altered white matter integrity in adolescents with prelingual deafness: a high-resolution tract-based spatial statistics imaging study. AJNR Am J Neuroradiol 34:1264–1270

Mower GD, Caplan CJ, Christen WG, Duffy FH (1985) Dark rearing prolongs physiological but not anatomical plasticity of the cat visual cortex. J Comp Neurol 235:448–466. doi: 10.1002/cne.902350404

Murray MM, Lewkowicz DJ, Amedi A, Wallace MT (2016) Multisensory Processes: A Balancing Act across the Lifespan. Trends Neurosci xx:1–13. doi: 10.1016/j.tins.2016.05.003

Noppeney U (2007) The effects of visual deprivation on functional and structural organization of the human brain. Neurosci Biobehav Rev 31:1169–1180. doi: 10.1016/j.neubiorev.2007.04.012

Noppeney U, Friston KJ, Ashburner J, et al (2005) Early visual deprivation induces structural plasticity in gray and white matter [1]. Curr. Biol. 15

Pascual-Leone A, Amedi A, Fregni F, Merabet LB (2005) The plastic human brain cortex. Annu Rev Neurosci 28:377–401

Pascual-Leone A, Hamilton R (2001) The metamodal organization of the brain. In: Progress in Brain Research. pp 427–445

Price DJ, Kennedy H, Dehay C, et al (2006) The development of cortical connections. Eur. J. Neurosci. 23:910–920

Ptito M, Schneider FCG, Paulson OB, Kupers R (2008) Alterations of the visual pathways in congenital blindness. Exp Brain Res 187:41–49. doi: 10.1007/s00221-008-1273-4

Rauschecker JP (1995) Compensatory plasticity and sensory substitution in the cerebral cortex. Trends Neurosci. 18:36–43

Reich L, Szwed M, Cohen L, Amedi A (2011) A Ventral Visual Stream Reading Center Independent of Visual Experience. Curr Biol 21:363–368

Reid RC (2012) From Functional Architecture to Functional Connectomics. Neuron 75:209–217

Ricciardi E, Bonino D, Pellegrini S, Pietrini P (2014) Mind the blind brain to understand the sighted one! Is there a supramodal cortical functional architecture? Neurosci. Biobehav. Rev. 41:64–77

Rockland KS, Ojima H (2003) Multisensory convergence in calcarine visual areas in macaque monkey. In: International Journal of Psychophysiology. pp 19–26

Rossion B, Hanseeuw B, Dricot L (2012) Defining face perception areas in the human brain: A large-scale factorial fMRI face localizer analysis. Brain Cogn 79:138–157. doi: 10.1016/j.bandc.2012.01.001

Saint-Amour D, De Sanctis P, Molholm S, et al (2007) Seeing voices: High-density electrical mapping and source-analysis of the multisensory mismatch negativity evoked during the McGurk illusion. Neuropsychologia 45:587–597. doi: 10.1016/j.neuropsychologia.2006.03.036

Sampaio-Baptista C, Johansen-Berg H (2017) White Matter Plasticity in the Adult Brain. Neuron 96:1239–1251

Saur D, Kreher BW, Schnell S, et al (2008) Ventral and dorsal pathways for language. Proc Natl Acad Sci 105:18035–18040. doi: 10.1073/pnas.0805234105

Schall S, Kiebel S, Maess B, von Kriegstein K (2013) Early auditory sensory processing of voices is facilitated by visual mechanisms. Neuroimage 1–10

Schroeder C, Lakatos P (2009) Low-frequency neuronal oscillations as instruments of sensory selection. Trends Neurosci 32:1–16. doi: 10.1016/j.tins.2008.09.012. Low-frequency

Shibata DK (2007) Differences in brain structure in deaf persons on MR imaging studied with voxel-based morphometry. AJNR Am J Neuroradiol 28:243–249

Shiell MM, Champoux F, Zatorre RJ (2014) Reorganization of Auditory Cortex in Early-deaf People: Functional Connectivity and Relationship to Hearing Aid Use. J Cogn Neurosci 21:1–14

Shimony JS, Burton H, Epstein AA, et al (2006) Diffusion tensor imaging reveals white matter reorganization in early blind humans. Cereb Cortex 16:1653–1661. doi: 10.1093/cercor/bhj102

Shu N, Liu Y, Li J, et al (2009) Altered anatomical network in early blindness revealed by diffusion tensor tractography. PLoS One 4:. doi: 10.1371/journal.pone.0007228

Siegelbaum SA, Kandel ER (1991) Learning-related synaptic plasticity: LTP and LTD. Curr Opin Neurobiol 1:113–120. doi: 10.1016/0959-4388(91)90018-3

Smith KM, Mecoli MD, Altaye M, et al (2011) Morphometric differences in the Heschl’s gyrus of hearing impaired and normal hearing infants. Oxford University Press, Pediatric Neuroimaging Research Consortium, Cincinnati Children’s Research Foundation, Cincinnati, OH, USA.

Song S-K, Sun S-W, Ramsbottom MJ, et al (2002) Dysmyelination Revealed through MRI as Increased Radial (but Unchanged Axial) Diffusion of Water. Neuroimage 17:1429–1436. doi: 10.1006/nimg.2002.1267

Tomassini V, Jbabdi S, Klein JC, et al (2007) Diffusion-Weighted Imaging Tractography-Based Parcellation of the Human Lateral Premotor Cortex Identifies Dorsal and Ventral Subregions with Anatomical and Functional Specializations. J Neurosci 27:10259–10269. doi: 10.1523/JNEUROSCI.2144-07.2007

Van Ackeren MJ, Barbero FM, Mattioni S, et al (2018) Neuronal populations in the occipital cortex of the blind synchronize to the temporal dynamics of speech. Elife 7: doi: 10.7554/eLife.31640

van Wassenhove V, Grant KW, Poeppel D (2005) Visual speech speeds up the neural processing of auditory speech. Proc Natl Acad Sci 102:1181–1186. doi: 10.1073/pnas.0408949102

von Kriegstein K, Kleinschmidt A, Sterzer P, Giraud A-L (2005) Interaction of face and voice areas during speaker recognition. J Cogn Neurosci 17:367–76. doi: 10.1162/0898929053279577

Wheeler-Kingshott C, Alexander D, Schneider T, Cercignani M (2009) Measuring axial and radial diffusivities in the brain. In: Proceedings 17th Scientific Meeting, International Society for Magnetic Resonance in Medicine. p 3590

Wheeler-Kingshott CAM, Cercignani M (2009) About “axial” and “radial” diffusivities. Magn Reson Med 61:1255–1260. doi: 10.1002/mrm.21965

Whittingstall K, Bernier M, Houde JC, et al (2014) Structural network underlying visuospatial imagery in humans. Cortex 56:85–98. doi: 10.1016/j.cortex.2013.02.004

Willenbockel V, Sadr J, Fiset D, et al (2010) Controlling low-level image properties: the SHINE toolbox. Behav Res Methods 42:671–684. doi: 10.3758/brm.42.3.671

Zatorre RJ, Fields RD, Johansen-Berg H (2012) Plasticity in gray and white: Neuroimaging changes in brain structure during learning. Nat. Neurosci. 15:528–536

Zeng C, Yang Z, Shreve L, et al (2012) Somatosensory Projections to Cochlear Nucleus Are Upregulated after Unilateral Deafness. J Neurosci 32:15791–15801. doi: 10.1523/JNEUROSCI.2598-12.2012

